# Cell shape anisotropy and motility constrain self-organised feather pattern fidelity in birds

**DOI:** 10.1101/2021.01.22.427778

**Authors:** Camille Curantz, Richard Bailleul, Magdalena Hidalgo, Melina Durande, François Graner, Marie Manceau

## Abstract

Cellular self-organisation can emerge from stochastic fluctuations in properties of a developing tissue^1–3^. This mechanism explains the production of various motifs seen in nature^4–7^. However, events channelling its outcomes such that patterns are produced with reproducible precision key to fitness remain unexplored. Here, we compared the dynamic emergence of feather primordia arrays in poultry, finch, emu, ostrich and penguin embryos and correlated inter-species differences in pattern fidelity to the amplitude of dermal cell anisotropy in the un-patterned tissue. Using live imaging and *ex vivo* perturbations in these species, we showed that cell anisotropy optimises cell motility for sharp and precisely located primordia formation, and thus, proper pattern geometry. These results evidence a mechanism through which collective cellular properties of a developmental pattern system ensure stability in its self-organisation and contribute to its evolution.

## Main text

Most animals display visible patterns classified in symmetries, trees, spirals, meanders, waves, tessellations, cracks, stripes, and spots. These geometries arise in developing tissues through the acquisition of positional cues that trigger spatially restricted molecular and cellular response. One such pattern-forming mechanism is self-organisation, in which spatial arrangement spontaneously emerges from the amplification and stabilisation of small-scale stochastic fluctuations^1–3^. This process is typically robust to perturbations, but its outcomes are highly sensitive to changes in properties of the system^4,5^. The malleability of self-organisation explains extensive diversity in animal’s natural patterns^6,7^, but is hardly compatible with the meticulous precision at which they are produced in individuals of a species, a fidelity necessary to guarantee survival and reproductive success^8^. A pressing problem is thus to identify mechanisms through which fluctuations are channelled such that self-organised developmental systems achieve pattern fidelity. In models of self-organised behavioural patterns (e.g., snail aggregations^9^, army ants collective structures^10,11^), pattern fidelity stems both from the complexity of environmental parameters and the simplicity of behaviours at individual level. In developing tissues, emergent patterns of cellular self-organisation can be influenced by the redundancy and complexity of genetic networks controlling cell behaviour and/or by individual system properties such as molecular diffusion and cell-driven force generation and transmission^5,12,13^. However for lack of comprehensive work linking these properties to natural pattern variation, the nature of events shaping precision in tissue response –and therefore in pattern geometries, has remained a black box.

### Self-organisation of primordia in species-specific regions produces varying geometries

To address this question, we studied the self-organisation of typical hexagonal feather follicle arrays in the dorsal avian skin (Fig. 1a; ^14^). During development, the skin divides in regions of follicle-forming competence marked by a higher homogeneous expression of *β-catenin* transcripts and its protein located at the membrane of overlying epidermal cells (i.e., competence stage). Each follicle precursor, or primordium, arises through local condensation of cells in the dermis and epidermis (i.e., condensation stage^15–17^). Numerical and empirical work showed that this cellular process results from self-organisation driven by mechanical properties of the skin tissue and molecular factors acting through both reaction-diffusion and chemotaxis^17–20^. Primordia initiate a genetic program of feather production, mechanically-inducing translocation of the β-catenin protein in the nuclei of epidermal cells^16^ (i.e., differentiation stage; Fig. 1b). Because feather primordia geometries are typical to each bird group^17,21^, primordia self-organisation is likely constrained by processes ensuring species-specific pattern fidelity. To identify those, we first compared dynamics of primordia array emergence between avian species (Fig. 1c). In the zebra finch *Taeniopygia guttata*, part of the species-rich clade *Neoaves*, and in the domestic chicken *Gallus gallus* and Japanese quail *Coturnix japonica*, representing the *Galloanserae* clade, primordia emergence occurred along thin *β-catenin*-expressing segments located bilaterally to the midline in the zebra finch and Japanese quail and medially in the domestic chicken. They emerged as readily round shapes, visualised with restricted *β-catenin* expression and DAPI/phalloïdin stains marking cell nuclei and peripheral actin. Their distribution created a longitudinal primordia row, to which new rows were juxtaposed laterally in a sequential wave, as previously described^17,21^. When β-catenin translocated in first-formed primordia, at least 3 rows were visible, together forming the hexagonal motif. By contrast, in the emu *Dromaeis novaehollanidae* and the ostrich *Struthio camelus*, both part of the ancestral-most group *Paleognathae*, patterning occurred in two wide *β-catenin* expressing areas on each side of the dorsal midline. First primordia emerged visibly simultaneously in budding shapes distributed in wavy oblique lines. They progressively acquired rounder shapes, together producing a spatially random primordia array. Finally, in the Gentoo penguin *Pygoscelis papua*, flightless *Neoaves* bird, primordia formed in a tight mesh throughout a medial *β-catenin*-expressing area (Fig. 1d and Extended Data Fig. 1). Thus, avian species displayed differences in primordia geometries independent of the presence of a sequential wave and that form within species-specific patterning spaces (i.e., competent regions restricted to thin segments or wider areas). We quantified pattern fidelity by recording variability in spacing between adjacent primordia along first-formed rows in the domestic chicken, Japanese quail, and zebra finch, and across portions of competent areas in the emu, ostrich, and penguin (see methods and Extended Data Fig. 2). We found that primordia emerged and remained regularly distributed in initial segments/areas in the Japanese quail, zebra finch and penguin, while they evolved from irregularly to regularly distributed in the domestic chicken, and displayed irregular distribution throughout time in the emu and the ostrich (Fig. 1e). Strikingly, these differences were not correlated to variation in primordia size and density (Extended Data Fig. 3), suggesting these attributes are controlled independently from pattern fidelity. Together, these results evidenced inter-species variation in timely evolution and final outcomes of pattern fidelity in self-organising primordia arrays.

**Fig. 1:**
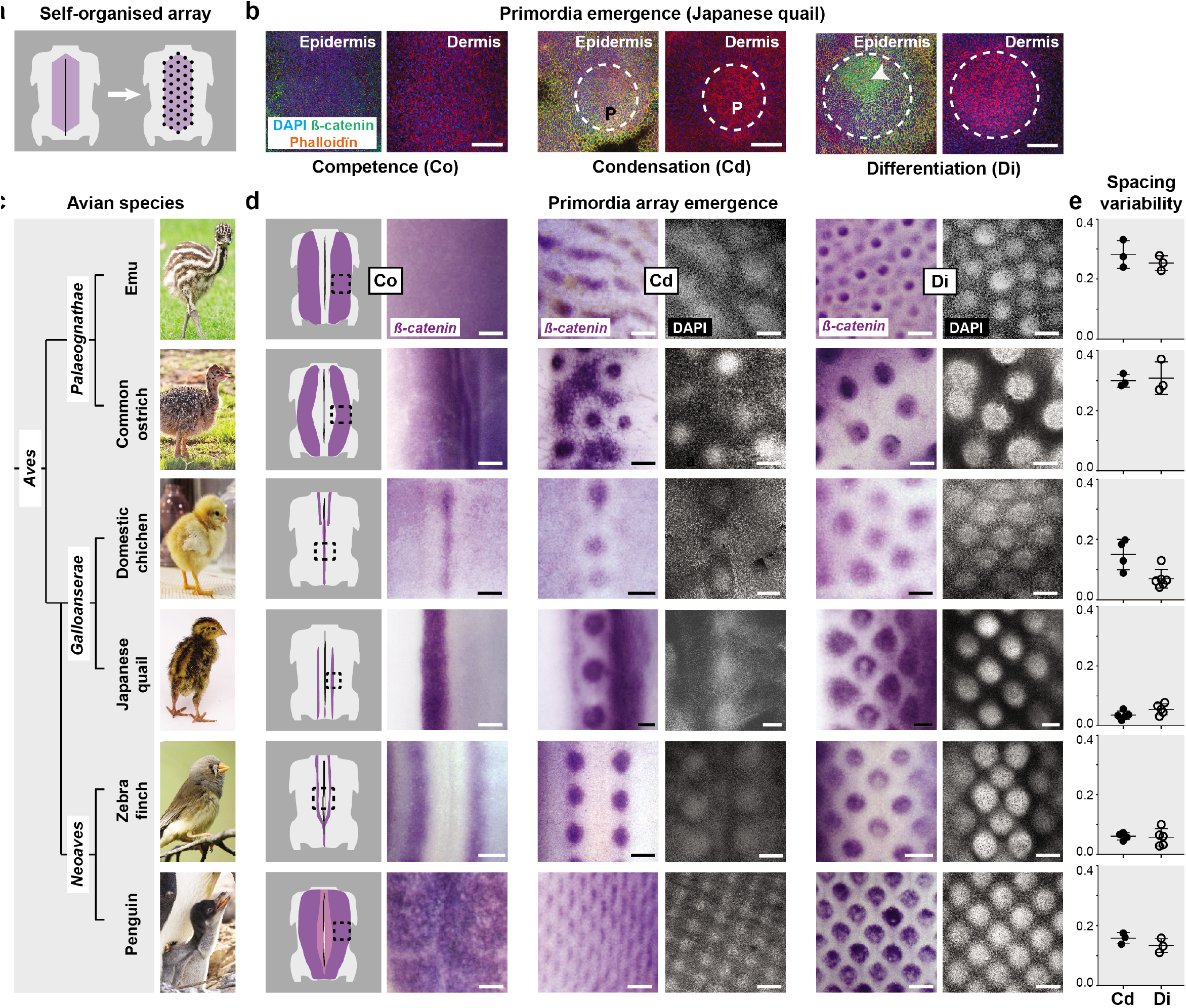
Natural variation in primordia array geometry between avian species. **(a)** Competent regions (in purple) in the dorsal skin of birds form primordia arrays (black dots) through cellular self-organisation. **(b)** 40X confocal views in competent regions of DAPI (in blue), β–catenin (in green) and phalloïdin (in red) stains in Japanese quail embryonic skins show that epidermal and dermal cells were homogeneously distributed prior to pattern formation (Competence stage Co) before compacting locally (Condensation stage Cd), forming nascent primordia (P, white dotted lines). Primordia initiate programs of feather production (Differentiation stage Di) upon nuclear translocation of β-catenin in epidermal nuclei (white arrows). Scale bars: 100μm **(c)** Juveniles of chosen bird species represent the three major clades of the avian phylogeny. **(d)** The earliest distribution of *β–catenin* transcripts (in purple) in competent longitudinal segments or areas at stage Co is shown on schematics of embryonic dorsum and images corresponding to black squares. At stage Cd, *β–catenin* was restricted to nascent primordia, also visualised with DAPI stains (in white, 10X confocal views), showing that species differ in timely dynamics of primordia emergence (i.e., in a row-by-row wave or simultaneously). At stage Di, species-specific primordia array geometries had formed. Scale bars: 200μm **(e)** Quantifying standard deviations of distances between adjacent primordia (spacing variability, in a dimensionless number defined in methods) along segments or areas at stages Cd and Di showed that species vary in the regularity of their primordia pattern. Error bars: mean with standard deviation. Photo credits: Jooin (emu), ©Barbara Fraatz (Pixabay, common ostrich), Publicdomain pictures (domestic chicken), Manceau laboratory (Japanese quail), Wikimedia (zebra finch), and ©Raphaël Sané (www.raphaelsane.com, Gentoo penguin).

### Early dermal cell anisotropy amplitude correlated with pattern fidelity

To identify emergent properties shaping pattern differences, we compared the density and shape of dermal and epidermal cells prior to, and during, primordia formation. We found that local cell density is typical to each species, not linked to position in the phylogeny, and evolving through time without following species-specific primordia spacing variability or final pattern geometry (Extended Data Fig. 4). This is consistent with previous work showing that cell density thresholds above which primordia emerge vary between birds^16,21^. Using phalloïdin stains and a custom-made image quantification software based on Fourier transform without segmentation of cell contours^22^, we quantified the anisotropy of average cell shape, defined here as the amplitude of deformation of skin cells and their angle with respect to the midline of the embryo (see methods). In the epidermis, cells were slightly elongated along the antero-posterior axis throughout the patterning process. Their anisotropy amplitude values did not correlate with pattern geometry at any stage (Extended Data Fig. 5). In the dermis of Japanese quail and zebra finch embryos, species in which primordia arise and remain regularly spaced, we also observed antero-posterior anisotropy in the competent segment prior to primordia emergence. Amplitude values remained higher in inter-primordia spaces at condensation stage, before dropping at differentiation stage. In the domestic chicken, in which primordia spacing progressively changes from irregular to regular, dermal cell anisotropy was first low, then transiently increased antero-posteriorly in inter-primordia spaces. In the emu and the ostrich, species with irregular patterns, dermal cells were isotropic at all stages. Finally, in the penguin, which displays immediate pattern regularity, dermal cells had strongest anisotropy values, maintained in the inter-primordia space during late primordia differentiation. Strikingly, penguin dermal cells were elongated along the dorso-ventral axis, orthogonally to those of poultry birds and the zebra finch (Fig. 2a and Extended Data Fig. 6). Differences in anisotropy orientation between species with a regular pattern correlated with the shape of the patterning space: restricted along the antero-posterior axis in the Japanese quail and the zebra finch, but not in penguins. Dermal cell anisotropy amplitude correlated through time with emerging and final primordia pattern fidelity (Fig. 2b) but not with primordia size or density (Extended Data Fig. 7). Together, these results suggested that high cell anisotropy amplitude is necessary to shape cell response such that precise, high fidelity primordia arrays, characterised by low spacing variability, emerge from self-organisation of primordia. In this scenario, dorso-ventral cell orientation in the penguin could compensate the absence of timely wave in a wide patterning space.

**Fig. 2:**
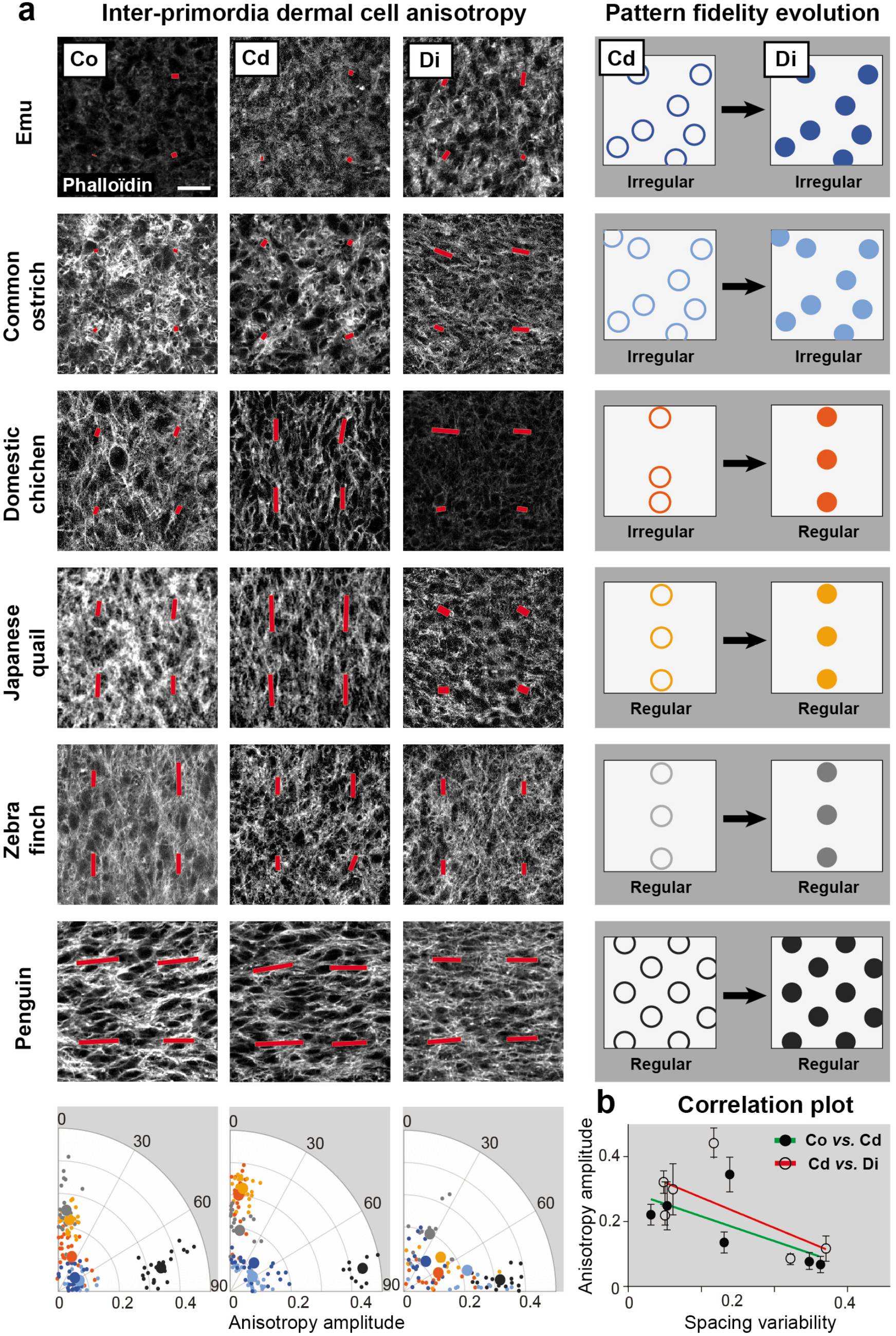
Dermal cell anisotropy amplitude correlates with primordia array geometry variation. **(a)** Left panels show confocal views at 40X of 100μm^2^ magnifications of phalloïdin-stained inter-primordia regions in the embryonic dermis of each species at stages Co, Cd and Di. Primordia emergence involves dynamic changes in the anisotropy of average cell shapes, which we extracted using Fourier transform and represented by red bars (^22^ and see methods; bar length is equal to measured anisotropy and bar direction is that of cell elongation). Right panel schemes represent the dynamic evolution of pattern array fidelity from stages Cd to Di in segments/areas for each species (see Fig. 1). Bottom panels show quantifications of anisotropy amplitude values (in a dimensionless number defined in methods) in all species, color-coded according to schemes and represented into polar coordinates for each stage (small dots are individual values, large dots are averaged values; n=3 specimen per species). **(b)** The plot shows correlating averaged values of dermal cell anisotropy at stage Co *vs*. spacing variability at stage Cd (black dots and green line; Pearson’s correlation coefficient r=-0.6769) or at Cd *vs*. Di (circles and red line; r=-0.6689). Error bars: standard deviation.

### Early dermal cell anisotropy is necessary for high pattern fidelity

To test these hypotheses, we perturbed cell anisotropy in the dermis prior to pattern emergence. We cultured competent dorsal skin portions of emus, which display dermal cell isotropy and pattern irregularity, Japanese quails, which have antero-posterior anisotropy and regular pattern, and African penguins *Spheniscus demersus*, closely-related to Gentoo penguins, and displaying identical dorso-ventral anisotropy and high fidelity pattern (Extended Data Fig. 8). Primordia emerged and underwent differentiation on cultured explants with proper timely dynamics though overall skin growth was impaired. Primordia density was higher (except in the emu) and primordia size decreased only in the penguin. (Extended Data Fig. 9). Primordia arranged in correct geometries in emu and Japanese quail skin explants, but pattern fidelity was lost in penguin explants compared to *in vivo* conditions (Fig. 3a). Thus, intrinsic properties of the developing skin and/or its substrate ensure the maintenance of the pattern geometry in this species. We found that dermal cell anisotropy remained species-specific in emu and Japanese quail explants, but dorso-ventral elongation observed in penguin flat skins was lost at places in explants (Fig. 3b). These observations are consistent with a role of dermal cell anisotropy in constraining pattern fidelity. To confirm this link, we further perturbed anisotropy by treating skins explants with Latrunculin A, drug known to inhibit actin polymerisation^23^. At low doses, drug treatment did not affect local cell density and efficiently decreased dermal cell anisotropy amplitude. Primordia forming on drug-treated explants had sizes and densities identical to control explants, emerging through proper timely dynamics (i.e., row-by-row in the Japanese quail), confirming that cell anisotropy does not control patterning sequentiality (Extended Data Fig. 10). In Latrunculin A-treated emu explants, spacing variability remained high, similar to control explants, and its dermal cells isotropic. In the Japanese quail however, drug treatment significantly increased spacing variability, and decreased cell anisotropy. In the penguin, spacing variability was not further increased, but primordia displayed elliptic or elongated shapes, and dermal cell anisotropy was lower (Fig. 3c and Extended Data Fig. 10). These results demonstrated that dermal cell anisotropy is necessary for the emergence and maintenance of low spacing variability. A minimal amplitude value appears required to achieve high fidelity of primordium arrangement.

**Fig. 3:**
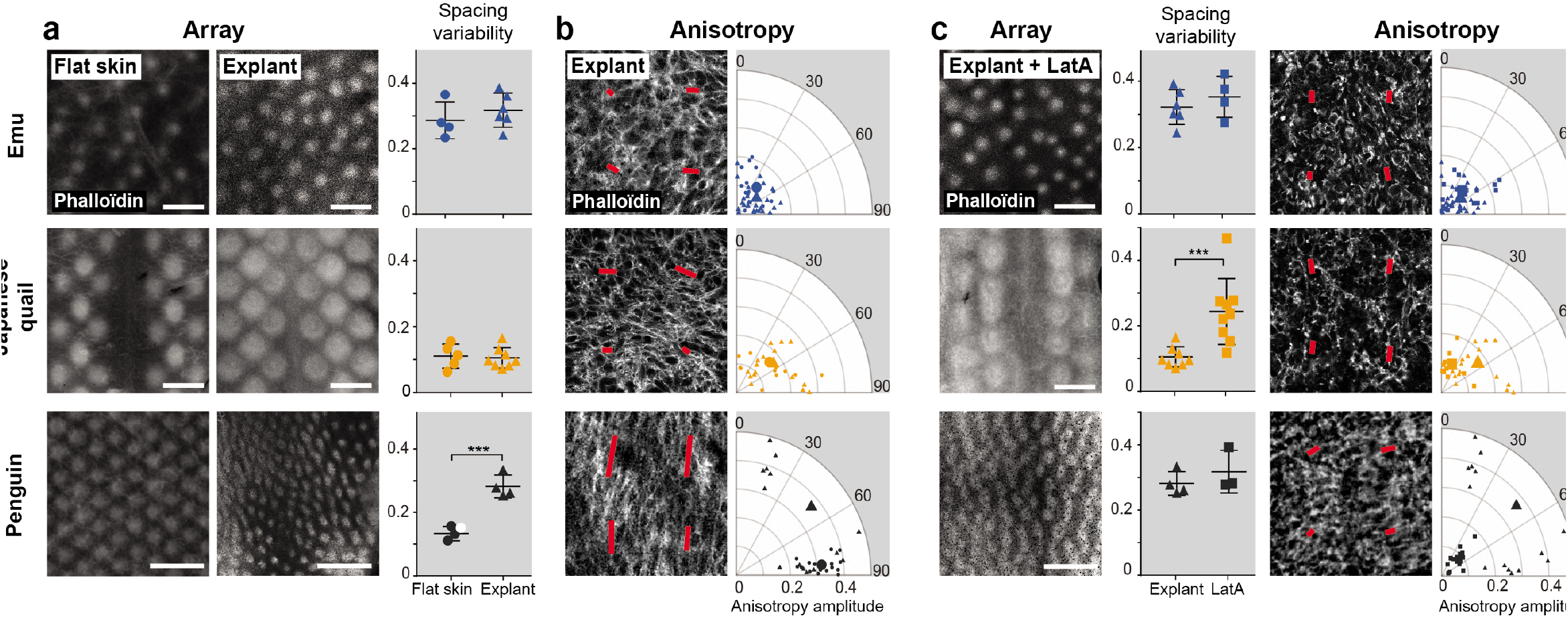
Dermal cell anisotropy amplitude constrains emerging pattern fidelity. **(a)** 3,2X views of phalloïdin-stained control flat skins (dots) and cultured explants (triangles) and quantifications of spacing variability in emus (n=5, in blue), Japanese quails (n=9, in orange) and penguins (n=3 for Gentoo penguins, in black; n=1 for African penguins, in white, see methods) show that at stage Di, pattern fidelity was maintained in emus (unpaired two-tailed *t*-test; p= 0.3963) and Japanese quails (p=0.7791) but significantly impaired in penguins (p=0.0005). Error bars: mean with standard deviation. **(b)** Confocal views and respective quantifications (as described in Fig. 2) of phalloïdin stained inter-primordia dermis of emu, Japanese quail, and penguin explants show that dorso-ventral anisotropy orientation at stage Di was lost in penguins compared to control flat skins shown in Fig. 2, cells at some locations orienting along the antero-posterior axis (as shown by large distribution of individual values, small triangles). **(c)** The same set of analyses performed in explants treated with low-doses of Latrunculin A (LatA, squares) showed that normal or culture-induced low pattern fidelity were maintained respectively in explants of emu (n=5; p=0.4195) and penguin (n=3; p=0.3928), and that high pattern fidelity was lost in explants of Japanese quail (n=9, p=0.002). In all species, cells were isotropic at this stage. Error bars: mean with standard deviation; significance of statistical tests is shown with stars.

### Dermal cell anisotropy modulates cell motility to sharpen primordia individualisation

To understand mechanisms linking early elongation of dermal cells to the production of high fidelity primordia arrays, we performed time-lapse imaging experiments encompassing the emergence of first primordia within competent skin segments of membrane-GFP (mb-GFP) Japanese quails (Fig. 4a and Supplementary video 1). We synchronised movies by using as reference time point *t_ref_* the visible condensation of dermal cells forming nascent primordia and applied to movies custom-made image analysis software for anisotropy detection. We found that continuous mean values of cell anisotropy amplitude provided sufficient criterion to automatically detect the competent region, and within this space, the putative primordia or inter-primordia regions. We analysed the dynamic evolution of dermal cell anisotropy in these software-defined areas and showed, consistent with results obtained on fixed tissues, that anisotropy was higher in the competent segment at least 4 hours before primordia emergence. Cell anisotropy then dropped progressively in the presumptive primordia region as local condensation became visible. We tracked the position of nascent primordia and found it remained stable through time, primordia being regularly spaced (Fig. 4b). Upon low-dose Latrunculin A treatment, which as expected led to a decrease in cell anisotropy values (Extended Data Fig. 11), nascent primordia often displayed transient elliptic shapes or smaller sizes, sometimes fusing together, events never occurring in control conditions (Supplementary Video 2). Tracking primordia position showed that compared to those of control explants, they were irregularly spaced and displayed higher overall displacement before merging and stabilising (Fig. 4c). These results showed that impairing cell anisotropy causes poor positioning of primordia through their increased displacement. We analysed cell movement using image correlation analysis (i.e., particle image velocimetry; see methods). Divergence analyses showed that Latrunculin A-treated explants undergo lower compaction in the forming primordia region compared to control explants, suggesting that observed transient elliptic shapes result from cell condensation defects (Extended Data Fig. 11). Cell motility remained stable between regions during primordia individualisation, its average speed consistent with values obtained in previous avian studies^21^ and data obtained in rodents^25^ but upon drug treatment, dermal cells were significantly more motile across the whole patterning space (Fig. 4d). These results contrasted with effects obtained with high doses of Latrunculin A, which block cell movement^21,24^. They demonstrated that a reduction of cell elongation correlates with freer dermal cell displacement, impairing proper primordia compaction and individualisation and lowering pattern fidelity.

**Fig. 4:**
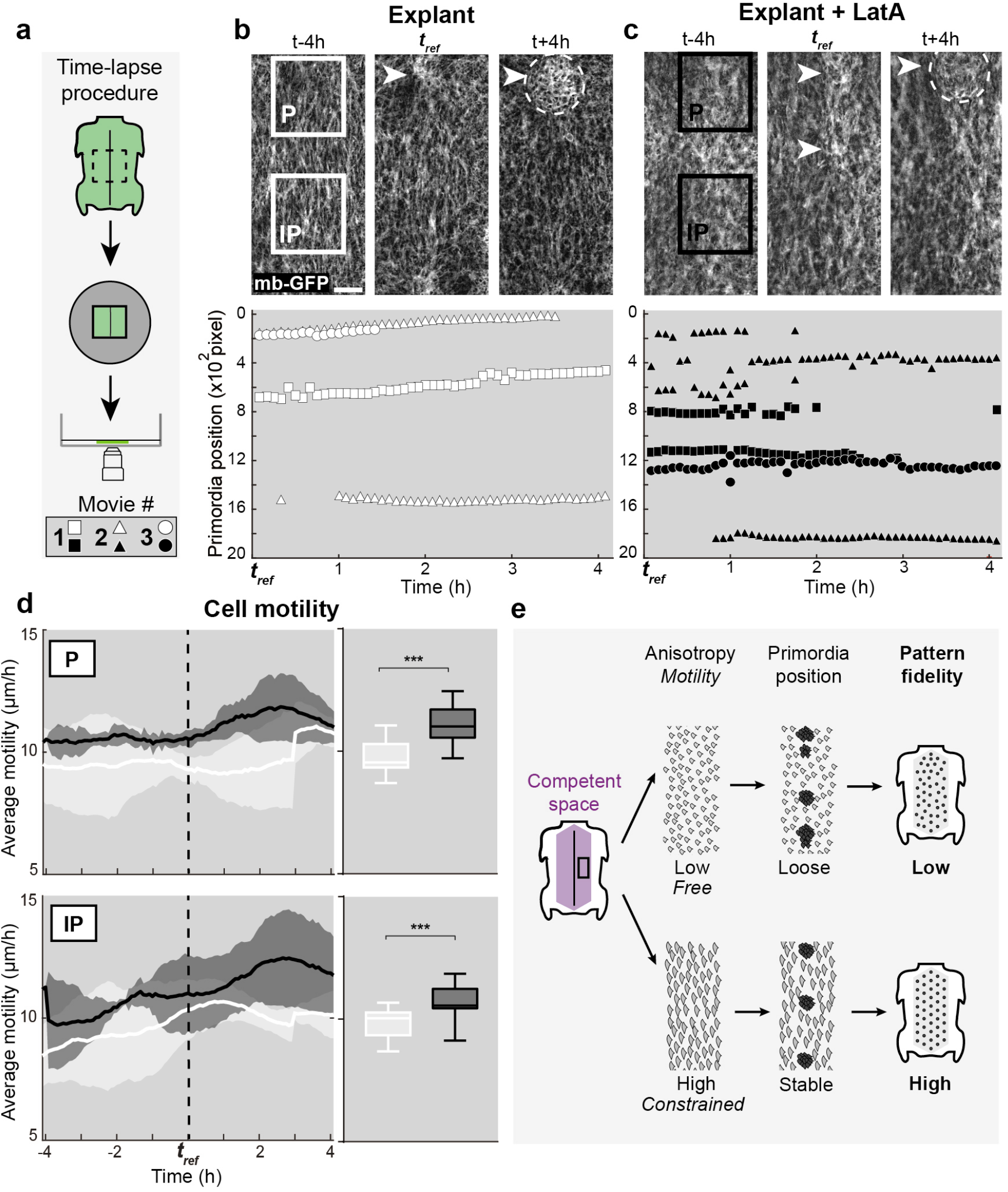
Anisotropy optimises dermal cell motility and nascent primordia positioning. **(a)** Schematic drawing of the time-lapse experimental procedure: dorsal skin explants were placed dermal side down on nitrocellulose filters for confocal imaging (see methods). Movie number is represented by a shape code. **(b)** Snapshots of time-lapse confocal movies 4 hours before, at, and after *t_ref_* recorded at dermal level on skin explants of membrane-GFP (mb-GFP) Japanese quails show the formation of a primordium (white arrow). White squared lines mark the position of automatically detected putative primordium (P) and inter-primordium (IP) areas. Tracking the antero-posterior position (in pixel x10^2^) of software-detected primordia at each time point of three independent movies (white shape-code as shown in the left panel) demonstrated that emerging primordia maintain stable positions. In movie#2, two spaced-out primordia emerge. Scale bar: 200μm. **(c)** The same experiments after Latrunculin A treatment (LatA; black squared lines mark P and IP areas in three movies (black shape-code) showed that reducing anisotropy causes imprecision in nascent primordia position, which can emerge closely and fuse together (movie#2 and #3). **(d)** Quantifications of average cell motility (in μm/h) through time (in hours to *t_ref_*, marked with a dotted line or averaged in box plots) in IP and P regions showed that cell motility increases upon drug treatment (in black) compared to control conditions (in white; unpaired two-tailed *t*-test, p<0.0001 both for IP and P). Error bars: standard deviation, stars show significance of statistical tests. **(e)** We uncovered a mechanism of pattern fidelity establishment through which early anisotropy constrains motility in dermal cells. This stabilises the position of nascent primordia and forms geometrically precise feather arrays.

This study evidences a mechanism through which self-organised pattern fidelity is achieved: early cell shape anisotropy, emerging property of the patterning space competent to form primordia, modulates cell motility –likely by imposing constrains on molecular and cellular interaction. High values of cell anisotropy thereby control local cell condensation, limiting tissue response to primordia self-organisation such that it sharpens their individualisation and stabilises their positioning, resulting in a high fidelity pattern (Fig. 4e). Thus, collective cellular properties are key to ecologically relevant canalisation of self-organisation. Our comparative approach allows suggesting they shape feather pattern geometry evolution. In low pattern fidelity flightless *Paleognathae*, delayed competence acquisition in response to molecular signalling^21^ is likely due to lower cell anisotropy and freer cell movement. In the alien case of penguins, extreme anisotropy allows achieving highest feather pattern regularity, key to water resistance, in the absence of spatial restriction of the patterning space. Cell shape changes may constitute an evolutionary novelty in this *Neoaves* bird adapted to aquatic life and extreme weather conditions. Further work taking advantage of such pattern variation will allow exploring initial conditions necessary to establish early anisotropy, including the role of previously implicated^16^ tissue mechanics.

## Materials and Methods

### Embryo sampling

Japanese quail, domestic chicken, emu and common ostrich fertilised eggs were obtained from local suppliers, respectively Cailles de Chanteloup, Les Bruyères Elevage, Emeu d’Uriage, and Autruche de Laurette / Autruche du Père Louis / Autruche de la Saudraye. Zebra finch fertilised eggs were collected from a breeding colony at the Collège de France. Gentoo penguin eggs were harvested from natural breeding sites of Stevely Bay, Grave Cove and Weddell Island in the Western Falkland Islands. This allowed obtaining a large number of Gentoo penguin eggs specimens, and this species was thus used for descriptive analyses of pattern geometries, cell density, and cell anisotropy. African penguin eggs were provided by the La Palmyre Zoo. These specimens were available in low numbers but allowed controlled incubation and laboratory work; this species was thus used in explant assays. All animal work was performed in compliance with regulations for animal use and specimen collection of the French Government and the European Council. The welfare of the zebra finch breeding colony was guaranteed through regular care and visits approved by official and institutional agreement (Direction Départementale de la protection des populations and Collège de France, agreement C-75-05-12). Research licenses for Gentoo penguin specimen collection have been granted by the Environmental Planning Department of the Falkland Islands Government (#R26.2017; #R43.2018; #R36.2019).

### Flat skins and cultured explants preparation

Flat skins specimens were prepared as described^26^ after egg incubation in Brinsea Ovaeasy 190 incubators in the laboratory, embryo dissection, and fixation in 4% formaldehyde. Skin explants were produced by dissecting dorsal portions of embryonic skins at competence stage, namely HH32 in the emu, HH29 in the Japanese quail, and HH30 in the African penguin (see justification for using this species in the first paragraph). Explants were placed dermal side down on culture insert membranes (Falcon #353103, #08-771-20) over 800-to-1600μL DMEM solution supplemented with 2% FCS and 2% Penicillin/Streptomycin. For drug treatment, 0.0625μM/ml Latrunculin A (Sigma #428021) was added to the culture media. Japanese quail skins were incubated at 37°C with a 5% CO2 atmosphere (Thermo Scientific Midi 40). Explant were cultured at 37°C in 5% CO2 atmosphere (Thermo Scientific Midi 40), either during 24h for the Japanese quail (i.e., to condensation stage; n=8 for control explants; n=9 for drug treated explants), or during 72h for the African penguin and emu, and 48h for the Japanese quail (i.e., to differentiation stage; n=9, 5 and 3, respectively, for both control and drug treated explants).

### Imaging

Fixed flat skins and whole embryos were imaged using an AF-S Micro NIKKOR 60-mm f/2.8G ED macro-lens equipped with a D5300 camera (Nikon) and an MZ FLIII stereomicroscope (Leica) equipped with a DFC 450C camera (Leica). Confocal images were obtained using an inverted SP5 microscope (Leica) with 10X (dry) or 40X (immersed oil) objectives. For time-lapse imaging, dissected dorsal skin regions from membrane-GFP Japanese quails produced at the Pasteur Institute (Dr Gros laboratory) at HH29, which corresponds to competence stage Co, were placed dermal side down on Millipore-nitrocellulose filters (Sigma #HABP04700) and cultured in a Lumox dish (ø50mm, Sarstedt #11008) containing 5 ml of DMEM supplemented with 2% FCS and 2% Penicillin/Streptomycin (and when applicable, 0.0625μM/ml Latrunculin A) at 37°C in 5% CO2 atmosphere. Time-lapse images were acquired every 5 minutes up to 10 hours with a SPINNING DISK-W1 (Zeiss) equipped with a 25X (immersed oil) objective.

### Quantification of pattern attributes

To assess primordia density at condensation and differentiation stages in the emu (n=5 and 3, respectively), Common ostrich (n=4 and 3), and Gentoo penguin (n=3 and 3, see justification for using this species in the first paragraph), species in which primordia form an array, we quantified the number of primordia per 1mm^2^ squares. For the domestic chicken (n=4 and 6), Japanese quail (n=8 and 8) and zebra finch (n=5 and 4), species in which primordia arise in longitudinal lines, we quantified the number of primordia in 0.3mm x 1mm rectangles within the first formed row of feather primordia and normalised values per mm^2^. We quantified primordia size at condensation and differentiation stages by manually outlining DAPI-stained dermal condensations with the freehand selection tool in Fiji software and recording obtained area value with Fiji measurement tools. 3 primordia of the first row or spread out in the array were measured for 3 different embryos in each species. Regularity of primordia arrangement was quantified at condensation and differentiation stages in the domestic chicken (n=4 and 6, respectively), Japanese quail (n=5 and 5) and zebra finch (n=5 and 5) as the standard deviation of distances (in μm) between neighbouring primordium centres in the first formed row, recorded on imaged specimens using Fiji measurement software and normalised to the mean value of spacing (in μm). In the emu (n=3 and 3), Common ostrich (n=3 and 3) and Gentoo penguin (n=3 and 3), primordia centres were detected automatically using a custom Matlab program based on image threshold adjustments (Dotfinder;^17^). Rare detection errors were manually corrected and the MATLAB function ‘delaunay’ was applied to the resulting set of points. The Delaunay triangulation was used to extract the distance between neighbouring primordia constituting a triangulation edge (in pixel); for each skin we quantified pattern regularity by computing the standard deviation of these edge lengths (edges with a vertex positioned on the boundary were excluded from the analysis) normalised to their average (in pixel). This quantification method is illustrated in Extended Data Fig. 2.

### Expression analyses

*In situ* hybridisation experiments were performed as described previously^27^ using antisense riboprobes synthesised from vectors containing 881-bp, 501-bp, and 685-bp fragments of coding sequences for *β-catenin* of the Japanese quail (also used for detection of expression domestic chicken, emu and Common ostrich), zebra finch, and Gentoo penguin, respectively. Digoxigenin-labelled riboprobes were revealed with an anti-digoxigenin-AP antibody (1:2,000, Roche) and an NBT/BCIP (Promega) substrate. Sequences of *β-catenin* primers were: F, AGCTGACTTGATGGAGTTGGA and R, TCGTGATGGCCAAGAATTTC for the Japanese quail, F: TAGTTCAGCTTTTAGGCTCAGATG and R: CCTCGACAATTTCTTCCATACG for the zebra finch, and F, GAACATGGCAACCCAAGCTG and R, GCCTTCACGGTGATGTGAGA for the Gentoo penguin.

### Immunohistological stains

Embryonic flat skin specimens were fixed in 4% formaldehyde overnight at 4°C, rinsed, and stained using anti-β-catenin (Abcam ab2365; 1:100) and goat anti-rabbit Alexa 488 (Abcam 11008; 1:500) antibodies. Co-stains were performed using Phalloïdin coupled to Alexa 546 (Abcam A22283; 7:200) and DAPI (Southern Biotech). Flat skins were mounted in Fluoromount (Southern Biotech) on slides prior to imaging.

### Quantification of cell density and anisotropy in fixed tissues

To quantify cell density, we counted the number DAPI-stained cell nuclei within 3 squares of 100μm^2^ distributed along the antero-posterior axis on focal plans of confocal images at epidermal or dermal levels (n=3 per species). To extract coarse-grained cell anisotropy we used our published Fourier transform (FT) based program (available on GitHub^22^) on images segmented in 256×256 pixel interrogation boxes with 50% overlap, as follows: we applied to each box a multiplying function to avoid singularities in the FT due to boundaries and computed the FT using the MATLAB fast FT algorithm (fft2.m). Resulting Fourier space patterns were binarised by retaining only the 5% brightest pixels yielding filled ellipses with correct aspect ratio, and inertia matrices were computed and diagonalised. This allowed extracting pattern anisotropy defined by 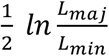 (where *L_maj_* and *L_min_* are respectively major and minor axes of the ellipse), a dimensionless number equal to zero when the pattern is isotropic and whose value corresponding to average elongation of cells within a given interrogation box is represented by the length and orientation of a bar (shown in red in Figures; see^22^ for further details). Primordia and inter-primordia regions were manually defined as groups of 4 interrogation boxes using stains mentioned above. Anisotropy amplitude was plotted as individual anisotropy bars for each group of 4 interrogation boxes (small shapes in polar plots); averaged anisotropy amplitude values were plotted as the mean of all bar lengths for a given species (large shapes in polar plots and correlation plots).

### Quantification of time-lapse imaging experiment

For time-lapse analyses, evolving values of anisotropy allowed defining regions automatically. Briefly, we computed average anisotropy for each interrogation box of each image throughout movies in which drift in “x” and “y” axes had been corrected using the “Stackreg” plugin in Fiji software. We defined the competent segment as the three adjacent columns with maximal averaged anisotropy. Within the herein defined competent segment, putative inter-primordia regions were the 4 interrogation boxes with maximal average anisotropy and primordia regions those with minimal average anisotropy. Anisotropy amplitude values in each region were first plotted separately for all movies. Resulting curves were smoothed using a moving average over 30 frames (i.e., 2h 30mins) and interpolated in the primordia region using MATLAB function “interp1” with evenly spaced query points. Movies of control and LatA-treated explants were compiled using as reference time point *t_ref_* the first visible condensation of cells in one primordium, also allowing computing and plotting dynamics of average anisotropy amplitude. For analyses of cell movement, we used MATLAB particle image velocimetry analysis program MatPIV on movies we treated with a 2 pixel radius median filter of Fiji software to eliminate aberrant vectors. We used reference time points *t_ref_* and averaging methods as above, primordia and inter-primordia regions were enlarged to 9 interrogations boxes (using the third column of the competent segment along the dorso-ventral axis and towards the other regions along the antero-posterior axis). The vector field obtained from MatPIV was used to quantify cell motility (averaging the norms of the 9 vectors in each region) and to assess divergent/convergent behaviours (using MATLAB function ‘divergence’ and averaging over the 9 vectors in each region). Tracking of primordia along the antero-posterior axis was carried out using the same threshold-based method as before (Dotfinder;^17^): for each image, we retained the 30% brightest pixels, applied Gaussian filtering (using Matlab function ‘imfilter’), binarised the image and detected primordia as white areas comprising at least 10000 pixels. All these methods are illustrated in Extended Data Fig. 11.

## Acknowledgements

We thank J. Gros and members of his laboratory for providing membrane-GFP Japanese quail fertilised eggs, as well as E. Bouvet and M. Ladner for their care of the zebra finch breeding colony. We thank C. Longtine for help with Gentoo penguin harvesting in the field, H. and M.P Delignières at Dunbar Farm and L. Clifton for providing landowner permission and accommodation for fieldwork at Grave Cove, Stevely Bay, and Weddell Island, and Denise Blake, Nick Rendell and the Falkland Islands Government for help with collection permits. We thank Ponant, Oceanwide Expeditions, all expedition staff and crew on-board, and A. Liddle for logistical help and support in assessing field sites. We thank T. Petit and the staff of the La Palmyre Zoo for their continuous help with supplying African penguin eggs. We thank the Collège de France imaging facility staff for help with time-lapse experiments, and S. Chanet for helpful comments on the manuscript.

## Author Contributions

CC collected specimens, quantified pattern attributes, cultured experiments, and time lapse imaging and analyses. CC, MH and RB performed *in situ* hybridisation experiments. RB and FG conceived quantitative computational analyses. MD wrote and implemented the software. CC and RB performed computational analyses. CC and MM conceived the project, designed experiments, and wrote the manuscript. This work was funded by an ERC Starting Grant (#639060) and a PSL University Grant to MM.

