## Extended data for "Cell shape anisotropy and motility constrain self-organised feather pattern fidelity in birds"

1 # Extended Data

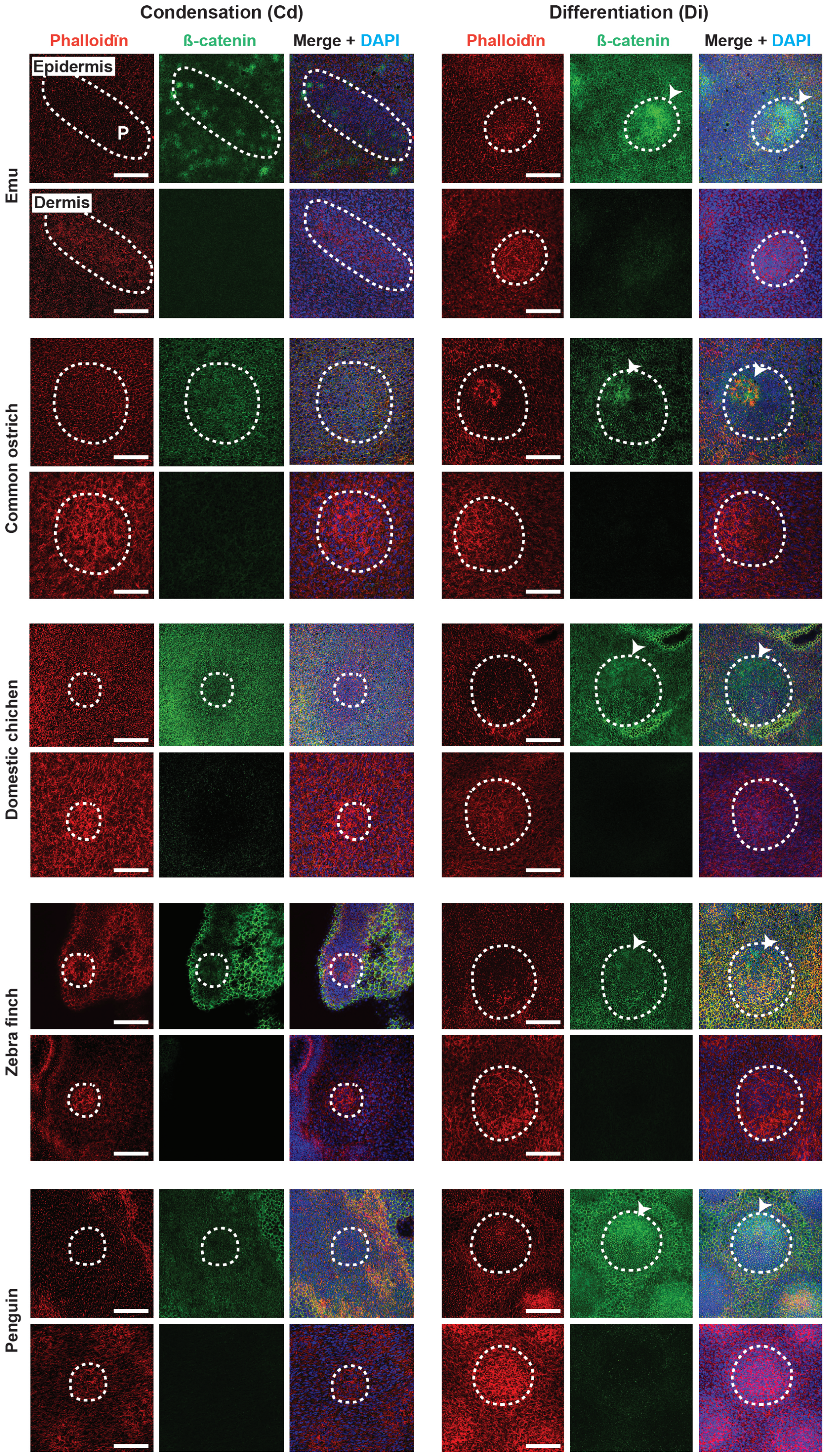


2

3

**Extended Data Fig. 1: Dynamics of primordia emergence in studied species.**

40X confocal views of DAPI (in blue), β–catenin (in green) and phalloïdin (in red) stains in flat embryonic skins of emu, common ostrich, domestic chicken, zebra finch and penguin show that similarly to the Japanese quail (see Fig. 1), epidermal and dermal cells compacted locally in primordia (P, white dotted lines) at condensation stage Cd and initiated programs of feather production upon nuclear translocation of β-catenin in epidermal nuclei (white arrows) at Differentiation stage Di. Scale bars: 100µm.

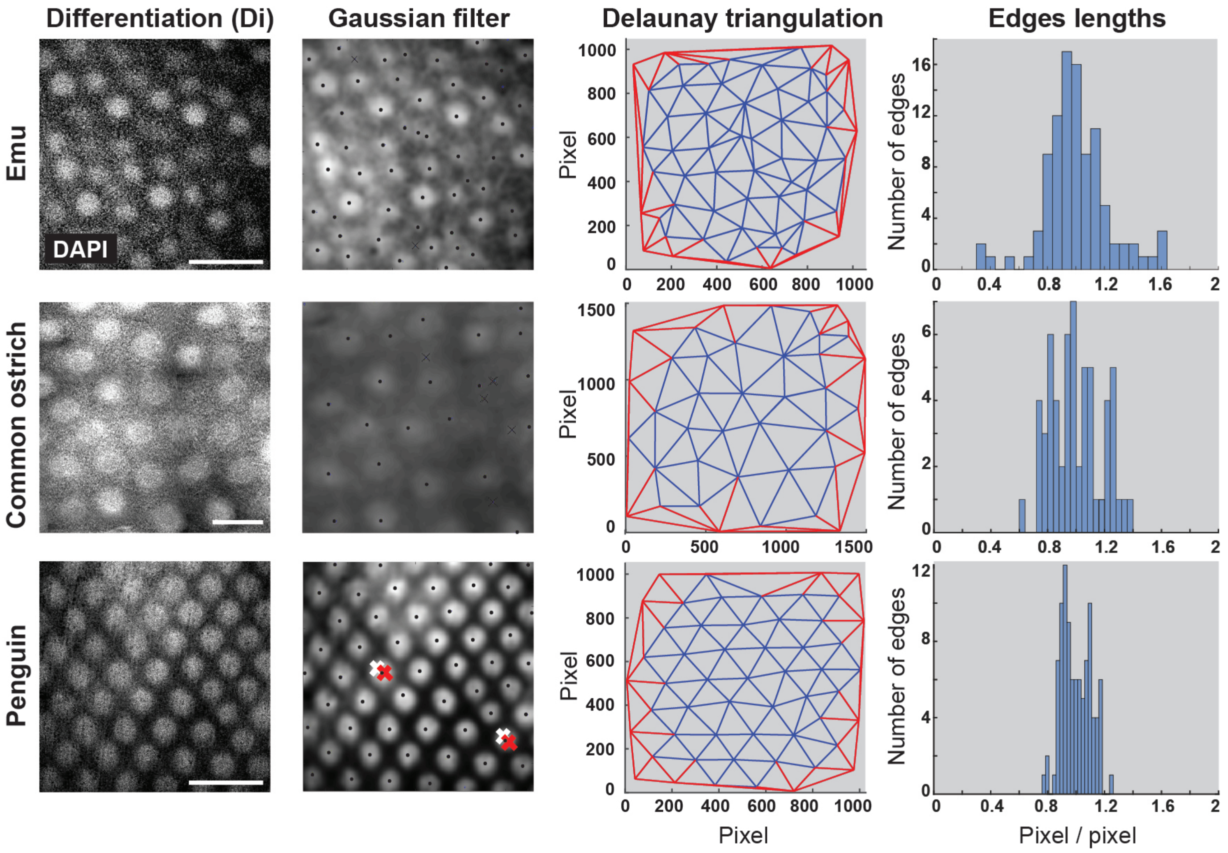


**Extended Data Fig. 2: Quantification of spacing variability in the emu, common ostrich and penguin.**

10X confocal views of DAPI stains (in white) in flat embryonic skins of emu, common ostrich and penguin at stage Di and associated positions of feather primordia centres (black dots) detected by applying a custom-made MATLAB program (Dotfinder;[17]) manually corrected in a few cases (crosses) allowed obtaining Delaunay triangulation representations. Triangles edges shown in red possess one vertex on the image boundary and were ignored in the analysis. Histograms show the distributions of edge lengths for each species (y-axis: number of edges) and illustrate spacing variability, quantified as standard deviation of normalised edge lengths (see methods and Fig. 2). Scale bar=500µm.

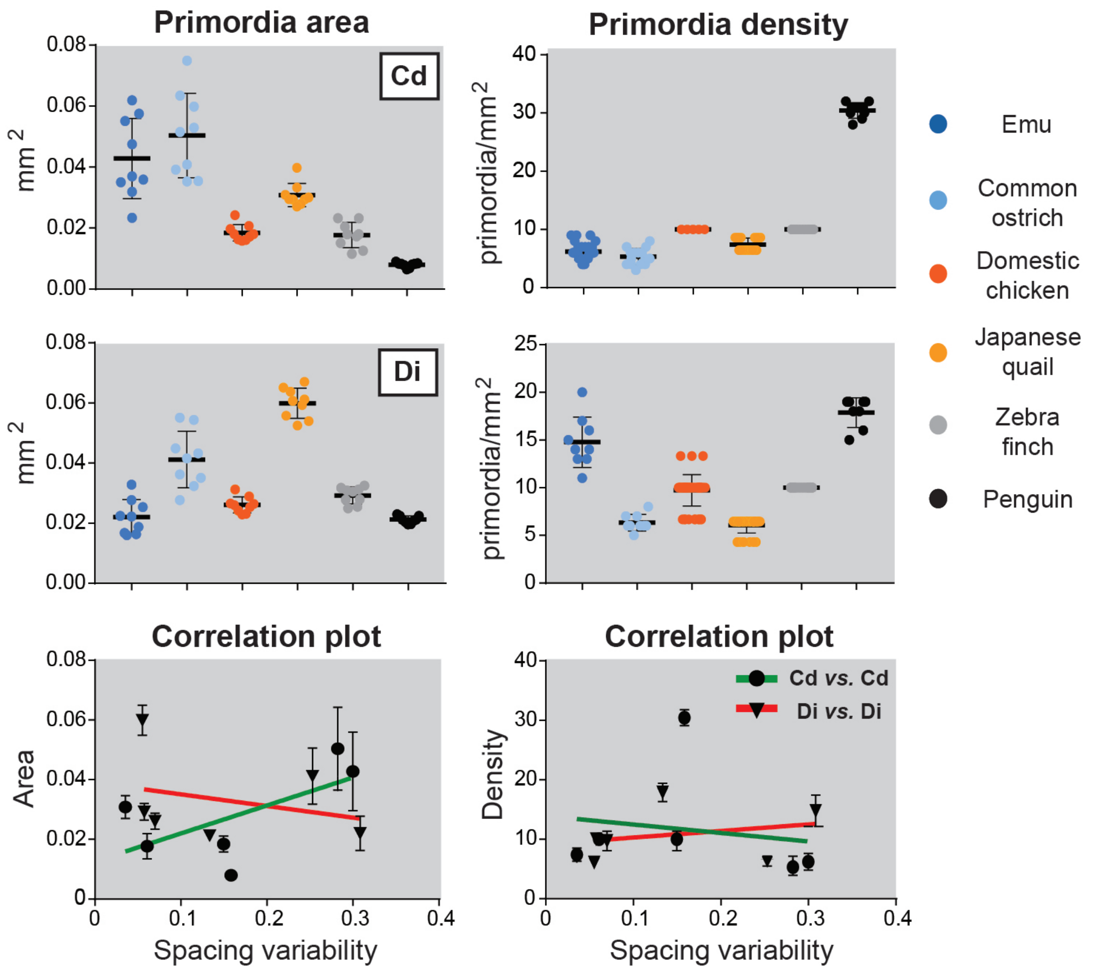


**Extended Data Fig. 3: Quantification of primordia size and density**

Quantifications of feather primordia size (in $mm^2$) and density (in primordia/$mm^2$) at stage Cd and Di are shown for each colour-coded each species. Plotting spacing variability values *vs.* average primordia area or density at stage Cd (dots and green line; Pearson's correlation coefficient r=0.6295 and 0.1654) and Di (triangles and red line; r=0.2865 and 0.2575) showed that spacing variability is not correlated to primordia size or density. Error bars: mean with standard deviation.

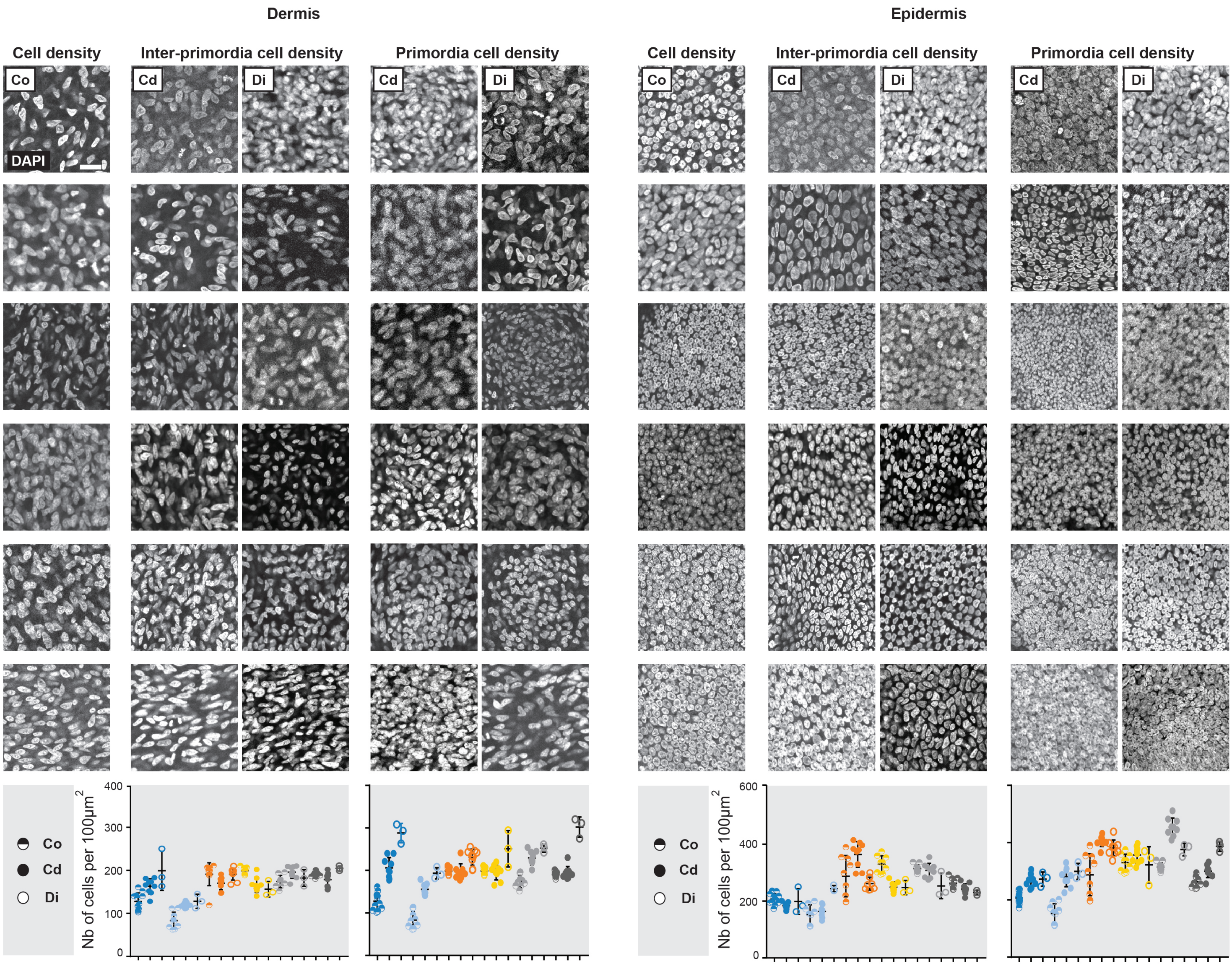


**Extended Data Fig. 4: Cell density at stage Co, Cd and Di in studied species**

40X confocal views at dermal and epidermal levels of DAPI stains (in white) in flat embryonic skins of emu, common ostrich, domestic chicken, Japanese quail and penguin at stages Co (bicoloured circles), Cd (black circles) and Di (white circles) in primordia and inter-primordia regions were used for quantifications of cell densities shown in corresponding graphs. Cell density increased through time, inter-species variation appearing largely independent of tissue level or stage. Scale bar: 20µm. Error bars: mean with standard deviation.

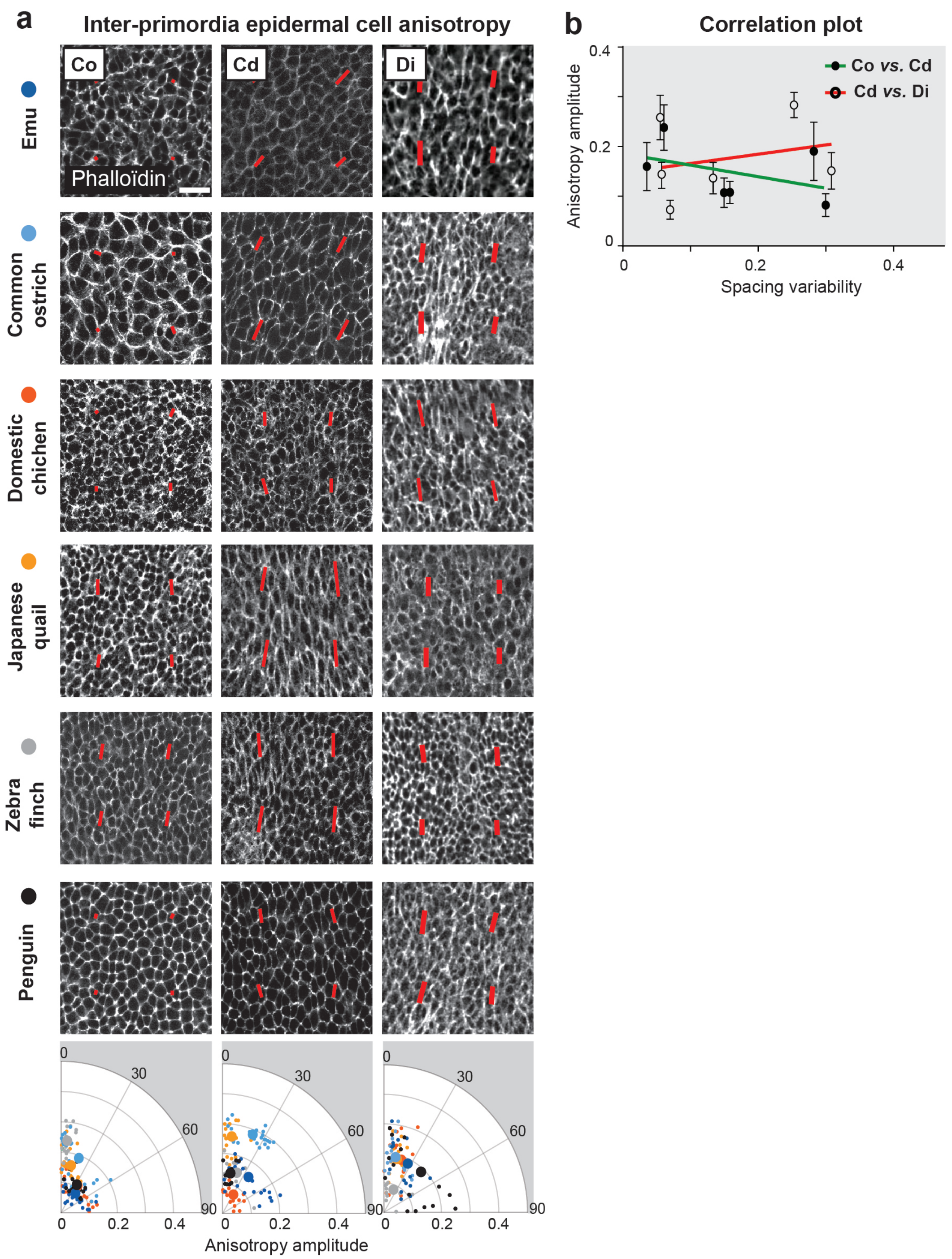

**Extended Data Fig. 5: Epidermal cell anisotropy in the inter-primordia region in studied species**

**(a)** 40X confocal views of 100µm$^2$ magnifications of phalloïdin stains (in white) in inter-primordia regions of the epidermis on flat embryonic skins of each species, at stages Co, Cd and Di, show dynamic changes in the anisotropy of average cell shapes (as described in Fig. 2; red bars). Quantifications of anisotropy amplitude in color-coded species are represented into polar coordinates for each stage (small dots are individual values, large dots are averaged values; n=3 specimen per species). Scale bar: 20µm. **(b)** The plot shows non-correlating values of averaged values of epidermal cell anisotropy at stage Co *vs* spacing variability at stage Cd (black dots and green line; Pearson's correlation coefficient r=-0.4274) or at Cd *vs.* Di (circles and red line; r=0.2543). Error bars: standard deviation.

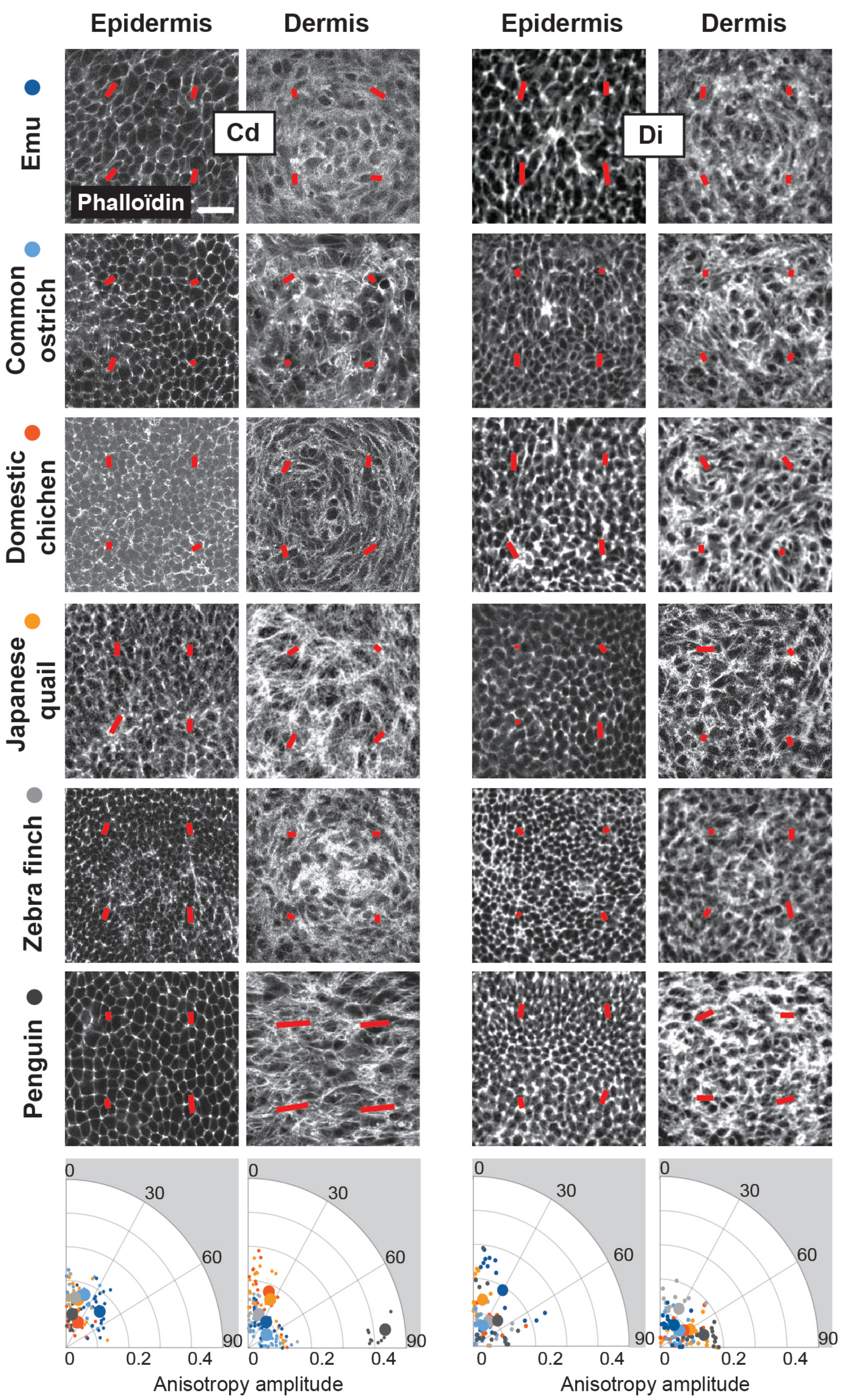


**Extended Data Fig. 6: Epidermal and dermal cell anisotropy in the primordia region in studied species**

40X confocal views of 100µm$^2$ magnifications of phalloïdin stains (in white) in primordia regions of flat embryonic skins in each species, at both dermal and epidermal levels, and at stage Cd and Di, show dynamic changes in the anisotropy of average cell shapes (as described in Fig. 2; red bars). Quantifications of anisotropy amplitude in color-coded species are represented into polar coordinates for each stage (small dots are individual values, large dots are averaged values; n=3 specimen per species). Scale bar: 20µm.

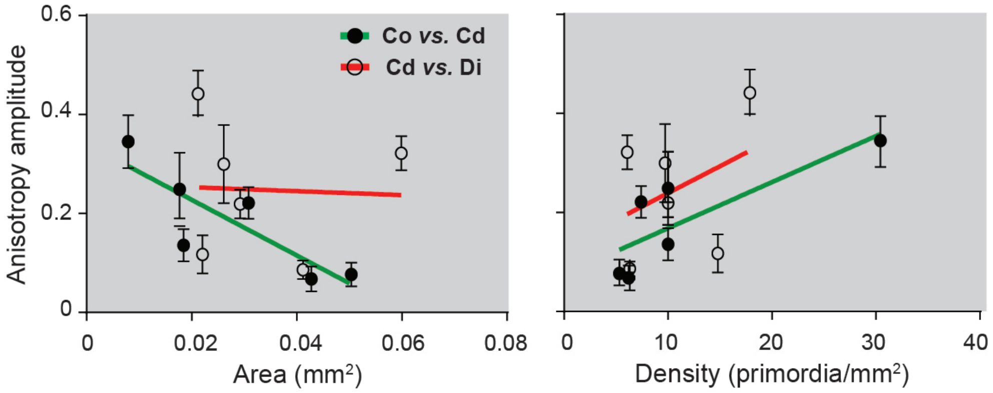


**Extended Data Fig. 7: Correlation of dermal anisotropy with primordia size and density**

Plots show non-correlating values of averaged dermal cell anisotropy amplitude at stage Co *vs.* primordia area or density at stage Cd (black dots and green line; Pearson's correlation coefficients r=-0.84305 or -0.8039), or at Cd *vs.* Di (circles and red line; r=0.04398 or 0.3800). Error bars: standard deviation.

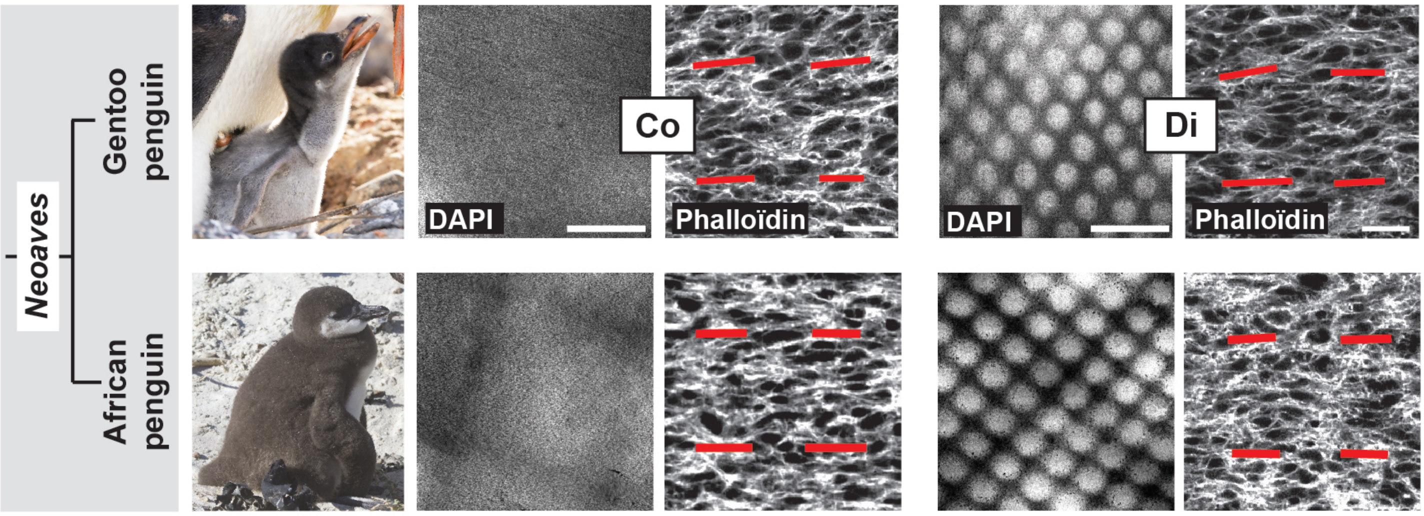


**Extended Data Fig. 8: Primordia array and dermal anisotropy in the African penguin**

10X confocal views of DAPI stains (left panels, scale bars: 500µm) and 100µm$^2$ magnifications of 40X confocal views of phalloïdin stains in inter-primordia regions (right panels, scale bars: 20µm) at stage Co and Di in embryonic flat skins of African penguin show that this species displayed pattern geometry and dermal cell anisotropy identical to those of the Gentoo penguin (see Fig. 2). Photo credits: ©Raphaël Sané (www.raphaelsane.com, Gentoo penguin) and ©Alain Bidart (www.alainbidart.fr, African penguin).

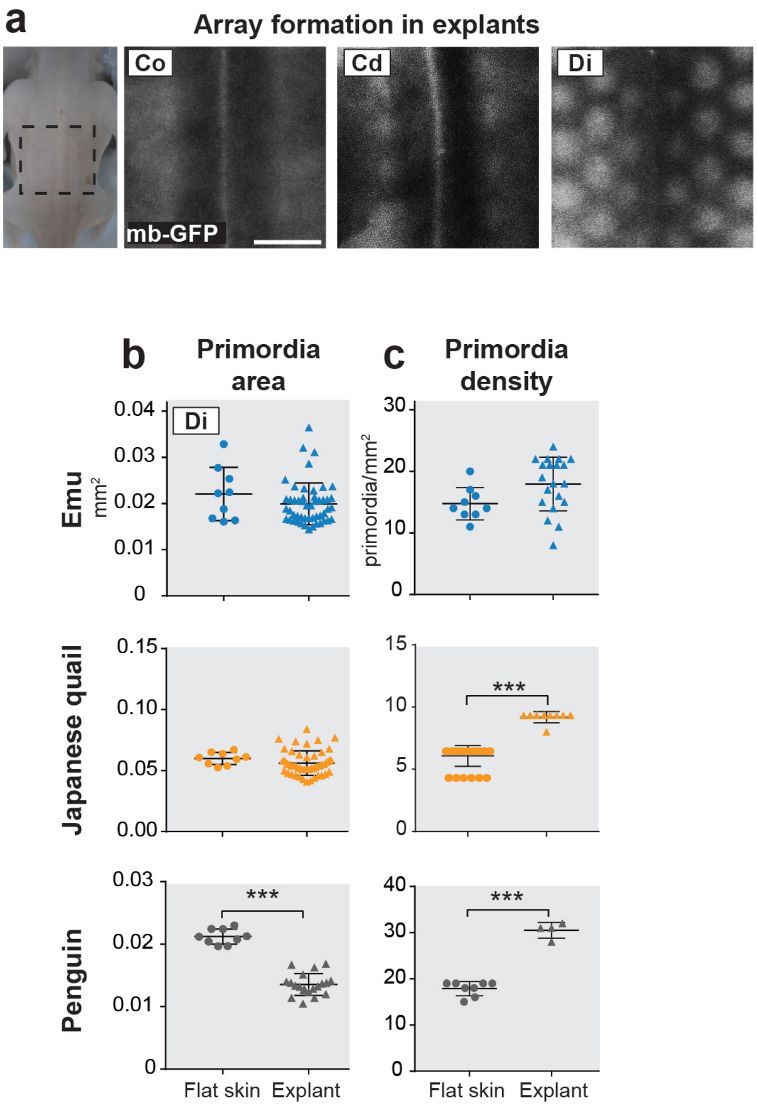


**Extended Data Fig. 9: Primordia size and density in cultured explants**

**(a)** In dorsal skin explants (corresponding to the dotted square) of transgenic membrane-GFP (mb-GFP) Japanese quail embryos cultured from stage Co to stage Di, primordia arrays forms with proper sequence. Scale bars: 500µm. **(b)** Quantifications of primordia size (in $mm^2$) at stage Di showed no significant difference between control flat skins and cultured explants in the emu (unpaired two-tailed *t*-test; p= 0.2180) and Japanese quail (p=0.2776), but a significant reduction in the penguin (p<0.0001). **(c)** Density (in primordia/$mm^2$) was conserved in the emu (p=0.0543) but significantly increased in the Japanese quail (p<0.0001) and penguin (p<0.0001). Error bars: mean with standard deviation; significance of statistical tests is shown with stars.

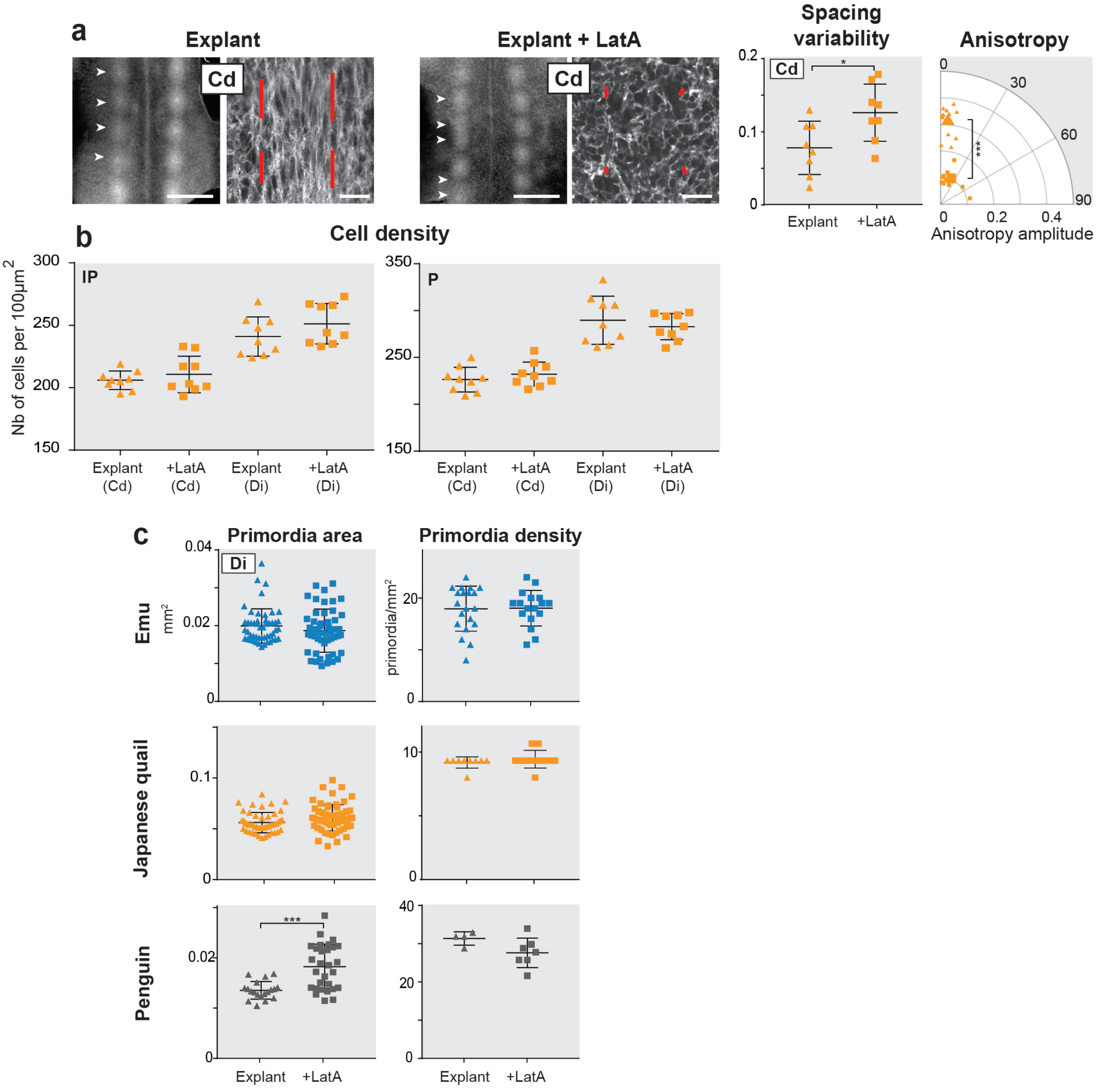


**Extended Data Fig. 10: Effect of Latrunculin on array geometry, cell density and anisotropy.**

**(a)** Phalloidin stained explants of Japanese quail at stage Cd are shown on confocal views at 10X (left panels, scale bars: 500µm) and 100µm$^2$ magnifications of 40X in inter-primordia regions (right panels, scale bars: 20µm). After Latruncilin A (LatA) treatment, antero-posterior anisotropy orientation was lost (red bars, see methods and Fig. 2). Quantifications of primordia spacing variability at stage Cd in control explants (triangles; n=8) or after drug treatment (squares; n=9) showed that Latrunculin A causes a significant decrease in pattern fidelity (unpaired two-tailed *t*-test; $p=0.0233$). Quantifications of dermal anisotropy amplitude represented into polar coordinates (small shapes are individual values, large shapes are averaged values; n=3 specimen) showed that Latrunculin A treatment significantly reduced anisotropy amplitude ($p<0.0001$). **(b)** Quantifications of dermal cell density normalised to $100\mu m^2$ areas at 3 different positions along the first formed row at stages Cd and Di showed no significant change between control (n=3) and drug-treated (n=3) Japanese quail explants in the inter-primordia region (IP; unpaired two-tailed *t*-test, p= 0.4076 at Cd and 0.1927 at Di) and primordia (P; p=0.3758 at Cd and 0.4895 at Di). **(c)** Primordia size (in $mm^2$) did not change between control and drug treated explants at stage Di of emus and Japanese quail (unpaired two-tailed *t*-tests; p=0.2363 and 0.1910) but was significantly increased in drug-treated African penguin explants ($p<0.0001$). Primordia density (in primordia/$mm^2$) did not change between control and drug treated explants in the three species (p=0.9341, 0.3369 and p=0.1054). Error bars: mean with standard deviation; significance of statistical tests is shown with stars.

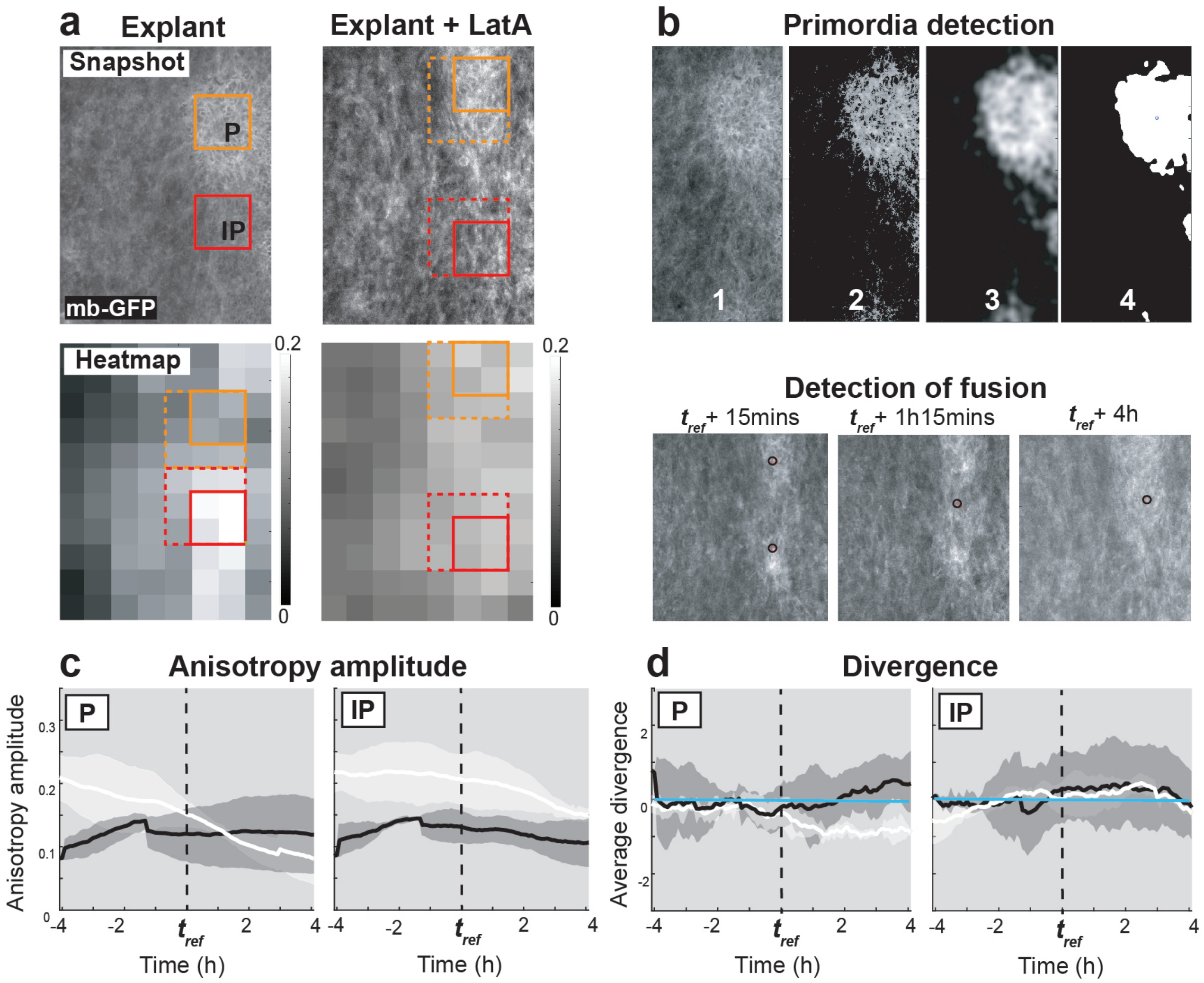


**Extended Data Fig. 11: Quantification methods for cell anisotropy, divergence and primordia tracking**

**(a)** Snapshots of time-lapse confocal movies on control (n=3) or Latrunculin A (LatA) treated (n=3) skin explants of membrane GFP (mb-GFP) Japanese quails were computed to produce heat-maps of average cell anisotropy amplitude per interrogation box (shades-of-grey squares). Over the course of each movie, putative inter-primordia (IP; red squares) and primordia (P; orange squares) regions were defined as the 4 interrogation boxes (for evolving anisotropy amplitude; full lines) or 9 interrogation boxes (for PIV analyses; dotted lines) with respectively maximal and minimal average anisotropy (see methods). **(b)** On time-lapse movie images (1) an algorithm detecting brightest pixels (2) then applying Gaussian smoothing (3) was used to automatically identify putative primordia (4) and track their positions along the antero-posterior axis through time (see Fig. 4b). In bottom panels, snapshots of a time-lapse movie, show that the algorithm first automatically detected two primordia (black circles) at *tref*+15min, then only one at *tref*+1h15min and *tref*+4h, thereby evidencing a fusion event. **(c)** Quantifications of dermal cell anisotropy amplitude within automatically defined IP and P regions showed it significantly decreased through time (in hours to $t_{ref}$, marked with a black dotted line) after drug treatment (black lines) compared to control conditions (white lines; unpaired two-tailed *t*-test; $p<0.0001$ for both IP and P). **(d)** Quantifications of the divergence of the vector field of dermal cell movement averaged within IP and P (the blue line represents value 0 at which there is no contraction or extension) showed that contraction in the primordium region, occurring 2 hours after $t_{ref}$, decreases upon LatA treatment (black lines) compared to control explants (white lines).

**Supplementary information**

**Supplementary Video 1:** A 8 hours time-lapse confocal movie (movie #2 in Fig. 4) of a membrane-GFP Japanese quail dorsal skin showed the dynamic emergence of two primordia at $t_{ref}$.

**Supplementary Video 2**: A 8 hours time-lapse confocal movie (movie #1 in Fig. 4) of a Latrunculin A-treated membrane-GFP Japanese quail dorsal skin showed the dynamic emergence of two primordia at $t_{ref}$.
